# Population genomics of *Macrophomina* spp. reveals cryptic host specialization and evidence for meiotic recombination

**DOI:** 10.1101/2024.09.03.609457

**Authors:** K.K. Pennerman, P. Goldman, C.J. Dilla-Ermita, G. Ramos, J.H. Jaime, J. Lopez-Hernandez, J. Ramos, M. Aviles, C. Borrero, A.O. Gomez, J.M. Neal, M. Chilvers, V. Ortiz, E.H. Stukenbrock, G.H. Goldman, A. Mengitsu, H.D. Lopez-Nicora, G.O. Sacher, N. Vaghefi, L. Kiss, J.P. Benz, A.R. Machado, T.E. Seijo, N.A. Peres, F.N. Martin, J.C. Broome, K. Ivors, G. Cole, S. Knapp, D.J. McFarlane, S.W. Mattner, M. Gambardella, P.M. Henry

## Abstract

Knowledge of the factors structuring populations of pathogenic fungi is fundamental to disease management efforts and basic biology. High-quality short-read sequence data were obtained for 463 *Macrophomina* spp. isolates collected from 91 host plant species and soil in 23 countries. Analyses revealed high diversity, admixture, and equal mating type ratios suggesting on-going recombination. Although most tested isolates could asymptomatically colonize strawberry, only isolates from a single phylogroup caused disease. In addition to strawberry, evidence for host specialization was discovered for soybean, demonstrating this broad host range pathogen contains phylogroups with cryptic specialization. Geography × isolate genotype associations were weak, suggesting these species were frequently trafficked between regions. Re-analysis using genomic data supported current species boundaries, and new molecular markers were designed to specifically identify each species. Contrary to expectations, *M. phaseolina* should be considered a species with both specialist and generalist populations for which meiosis can increase genetic diversity.

## Introduction

Effective management of plant disease requires knowledge of the factors structuring pathogen genomic diversity, such as host and/or geographic specialization, and the mechanisms driving the evolution of new pathogenic strains. For example, host specialization can occur at the sub-specific level^1,2^, and it must be understood to develop accurate diagnostic methods and evaluate host resistance. Associations between geographic regions and pathogen genotype can illuminate routes of dispersal that could be disrupted by phytosanitary regulations^3–5^. Additionally, the frequency of meiotic or parasexual recombination can indicate the likelihood for new, virulent strains to emerge^6^. A durable understanding of these factors can only be achieved with globally representative sampling, as international trade continues to move pests and pathogens to new areas^7,8^. These factors are only characterized for a few well-studied fungi, and expanding research to novel systems is necessary to broaden knowledge of fungal evolution.

The genus *Macrophomina* contains economically important plant pathogens, such as the cosmopolitan type species, *M. phaseolina,* which was first identified in the early 1900s, has been reported worldwide^9^, and can cause total crop losses in soybean and strawberry^10,11^. As a species, *Macrophomina phaseolina* has a broad host range of at least 97 plant species in addition to humans with compromised immune systems^9^. It is widely speculated that abiotic stresses induced by climate change will lead to increased damage by *M. phaseolina*^12^, as this fungus causes greater disease severity in warmer and drier conditions^13–16^. Four other *Macrophomina* species were recently described: *M. pseudophaseolina*^17^, *M. euphorbiicola*^18^, *M. tecta*^19^, and *M. vaccinii*^20^. These species have only subtle differences in morphology and were probably mis-identified as *M. phaseolina* before the widespread adoption of molecular phylogenetics^21^. Thus, their global distribution and host range are unknown and could be much wider than is reported.

Efforts to describe host specialization in *M. phaseolina* have usually relied on comparing isolates from different hosts without knowledge of their genomic differences, and this approach has not yielded clear results^22–25^. An exception to this trend is for strawberry-associated strains of *M. phaseolina*, which were found to cause severe strawberry disease in Spain, Argentina, and California, USA^26–29^. However, the broad applicability of these findings is unknown, as no association between host isolation source and virulence in strawberry was found in Israel^30^, and the populations of *M. phaseolina* from strawberry in Chile and Spain appear to be genetically distinct^31^. High resolution genotypic information is needed to resolve these discrepancies and illuminate the status of host specialization in *M. phaseolina*.

While isolate genotype × geography associations have been reported for specific populations of *M. phaseolina*^31,32^, it remains unknown if the global population of *Macrophomina* spp. is structured by geography, or if specific genotypes are found in disparate locations. A geographic influence on population structure seems plausible, given the apparent absence of aerial dispersal and the fact that specific environmental conditions play a large role in disease development. However, there is also some evidence for the ability of *Macrophomina* spp. to asymptomatically infect plants^28,33–36^, and be seedborne^37–41^ so they could have a high potential for dissemination between geographic regions.

Continuous adaptation to host and environment would be accelerated by on-going meiotic recombination. Many fungi that were historically considered asexual^17,42^ are now known to undergo meiosis at least periodically^43^. Both mating types exist in *M. phaseolina*^44^, but their ratios and the extent of admixture are unknown in extant populations. Gene flow could also occur through horizontal transfer of giant *Starship* transposable elements, which carry accessory genes to recipients^45^. Parasexual recombination can happen during anastomosis of vegetatively compatible hyphae^46,47^, but allo-recognition systems usually prevent anastomosis between genetically distinct strains^46–48^. Because of this, parasexual recombination would be much less likely to produce novel genotypes than meiotic recombination, which could occur between diverse genotypes^49^. Exploring the potential for meiosis in *Macrophomina* spp. is key to understanding the evolutionary potential of this genus.

To fill these knowledge gaps, the objectives of this work were to evaluate the evidence for host specialization, geographic adaptation, and recombination using a global survey of *Macrophomina* isolates from diverse geographic, temporal, and host sources. This investigation is critical to the global efforts to control diseases caused by *Macrophomina* spp., which are increasing along with abiotic stresses induced by climate change, and for gaining a broad understanding of the factors shaping fungal evolution at a global scale. To our knowledge, this is the first global genomic diversity study for any genus or species in the Botryosphaeriaceae.

## Results and Discussion

### A global genomic survey revealed fourteen genotypic clusters of *Macrophomina* spp

Whole genome shotgun sequences were obtained for 463 *Macrophomina* spp. isolates (435 newly sequenced and 28 retrieved from NCBI GenBank; Supplementary Table S1). These isolates were derived from 23 countries, 91 plant hosts and soil. Collection years ranged between 1927 and 2021, and therefore included many isolates described as *M. phaseolina* by morphology alone and/or before before 2014, when additional species were described within the genus *Macrophomina*. The most common hosts in our study reflected the most frequently researched hosts of *Macrophomina* spp. identified by Pennerman et al.^9^ (Supplementary Figure S1).

High-quality, bi-allelic single nucleotide polymorphisms (SNPs) with presence in at least 90% of isolates were identified using reference genomes of strains: Mp11-12 (from strawberry^27^; 578,310 SNPs; average of 8.07 SNPs per kbp for scaffolds with SNP loci), AL-1 (from alfalfa^27^; 588,334 SNPs; 9.39 SNPs per kbp), and mp117 (from soybean field soil^45^; 584,184 SNPs; 10.32 SNPs per kbp). Using these SNPs, fourteen population clusters were identified by a combination of admixture models, reticulate phylogenetic trees and principal component analyses (PCAs; Figure 1; Supplementary Figures S2 and S3). The cluster assignments were largely congruent among references. The only discrepancies were related to cluster assignment of highly admixed isolates F285 (at least 10% membership by Q-value in clusters VIII, IX, XIII and XIV) and F385 (clusters II, III, IX and XI). Cluster assignments in figures throughout this work are shown using the Mp11-12 reference, unless otherwise noted. Two clusters contained isolates that were not described as *M. phaseolina*: cluster I contained *M. euphorbiicola* and *M. pseudophaseolina* (12 isolates), and cluster II contained eight *M. tecta*, one putative *M. vaccinii* isolate, and two *M. phaseolina* isolates that were highly admixed with *M. tecta* (11 isolates total). The remaining twelve clusters contained isolates described as *M. phaseolina*.

**Fig. 1.**
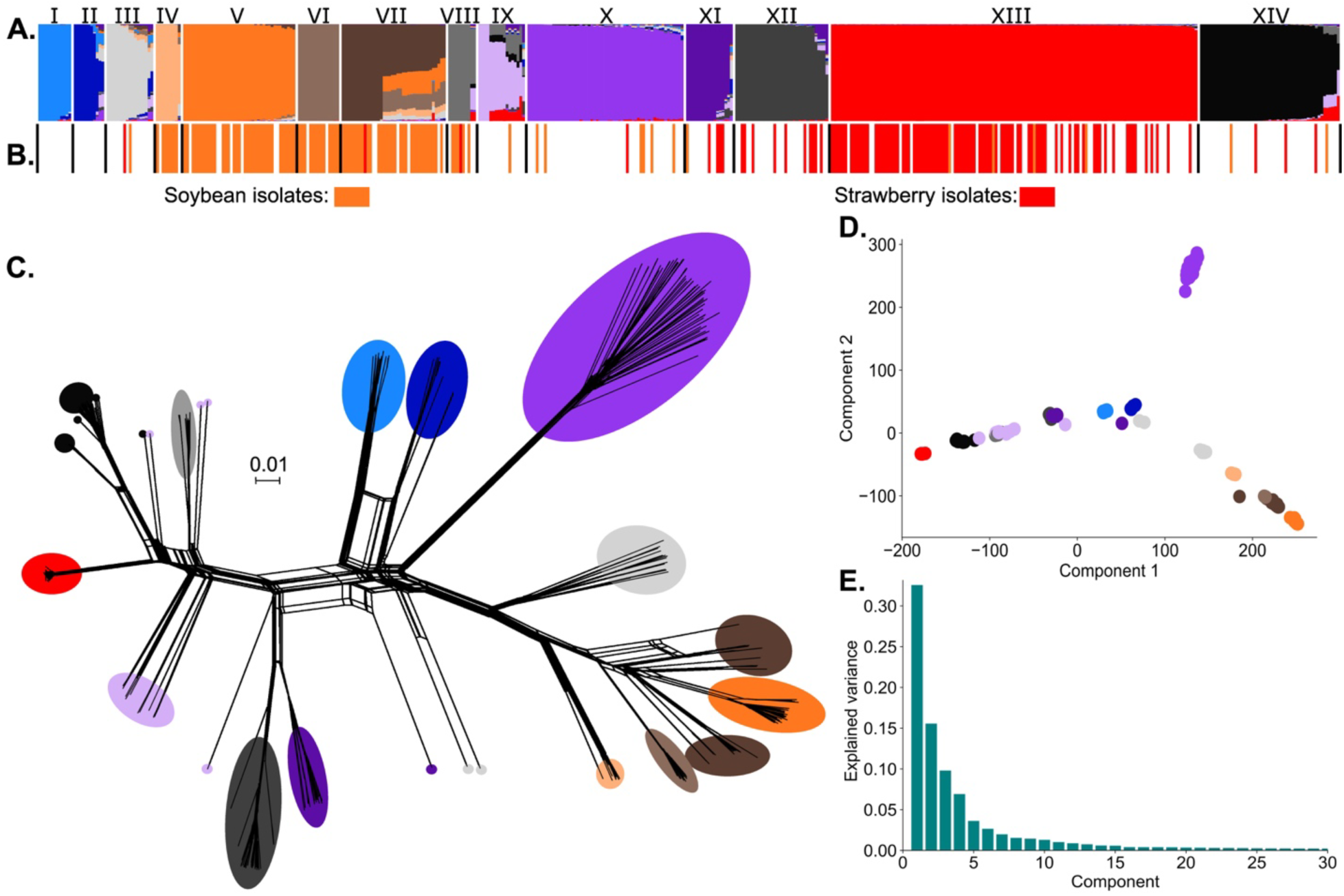
Grouping of *Macrophomina* spp. isolates with Mp11-12-based SNPs. The admixture plot (**A**) was generated with sNMF at K=14. From left to right, the clusters are labeled with Roman numerals and color-coded. (**B**) Colored bars indicate isolation source for each isolate in the above admixture plot, where strawberry = red, soybean or soybean field soil = orange, and cluster borders = black. The reticulate phylogenetic tree (**C**) and PCA plot (**D**) use the same colors as the admixture plot to indicate the major cluster assignment. The Scree plot (**E**) shows the eigenvalues for the first 30 principal components.

### Pangenomic gene content was structured by the 14 phylogroups

A pan-genomic analysis of gene content was conducted to evaluate the strength of the phylogroup designations and the genetic diversity within *Macrophomina* spp. A total of 113,290 homologs were identified across all genome assemblies. These homologs comprised 10,984 “super-core” homologs (present in all 463 *Macrophomina* spp. isolates), 18,866 “core” homologs (present in at least 95% of the isolates), and 45,233 “accessory” homologs (present in at least two isolates but fewer than 95% of all isolates; Figures 2A, 2B, 2C). Hierarchical clustering based on core and accessory homolog presence/absence variation yielded groups that were congruent with those identified by admixture and PCA analyses (Figure 2D). There were 38,207 singletons, and the pangenome appeared slightly open when these genes were included. When singletons were excluded, the pangenome structure was clearly closed (Figure 2E). The rarefaction observed for highly sampled clusters (e.g. V, VII, X, XII, XIII and XIV), suggested that continued sampling would have corroborated our finding of a closed pangenome structure (Figure 2F).

**Fig. 2.**
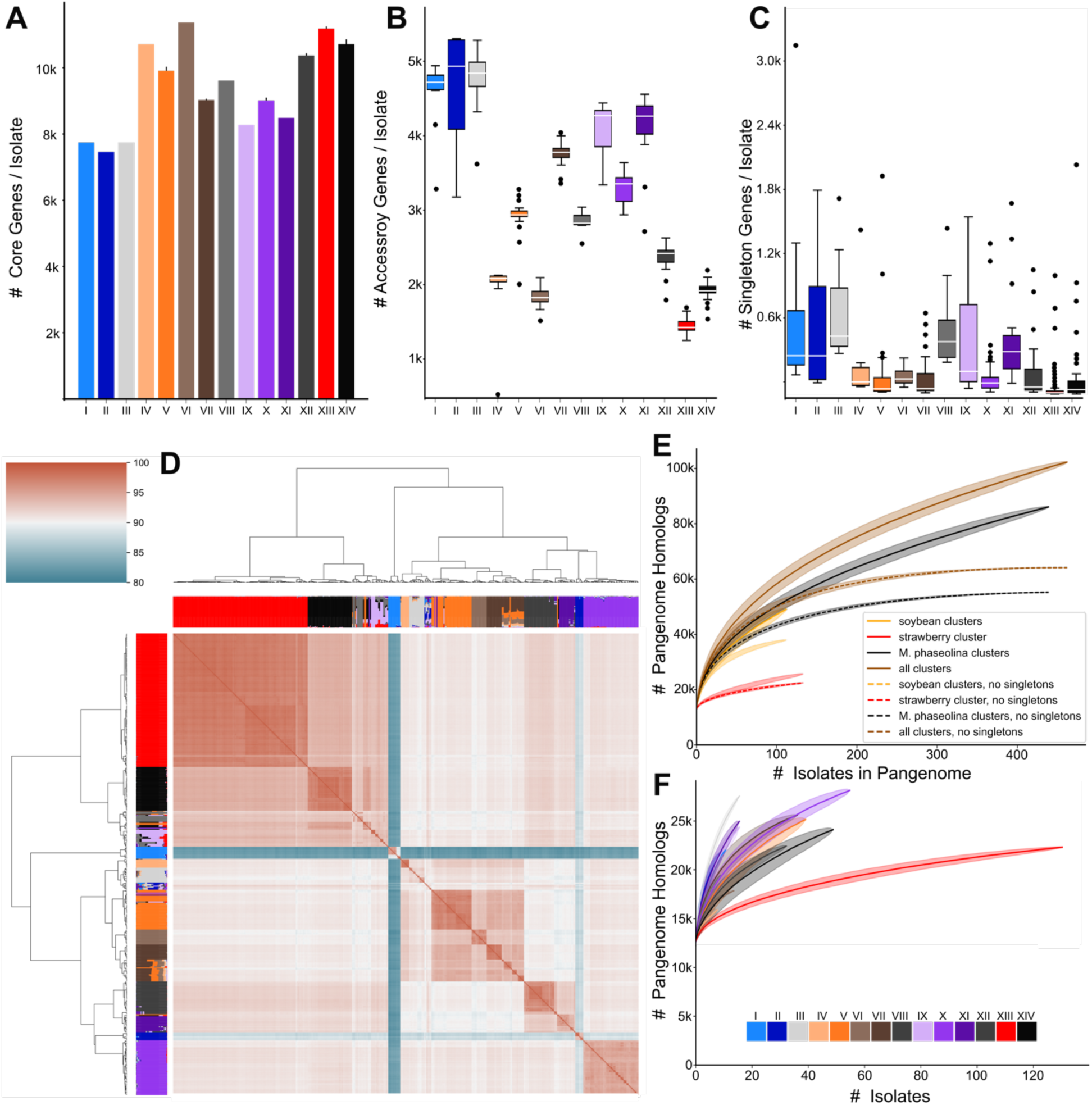
Homolog presence/absence variation by population cluster. **A.** Barplot depicting the average number of core gene homologs (present in >95% of isolates) for each population cluster. Error bars represent one standard deviation above and below the mean. **B.** Boxplot depicting the number of accessory gene homologs (present in more than 2 isolates, less than 95%) for each population cluster. **C.** The number of singletons (genes present in a single isolate) for isolates in each population cluster. **D.** A hierarchically clustered heatmap of core and accessory homolog presence/absence variation. Color intensity within the heatmap indicates the percent of shared homologs for each pair of isolates. The corresponding Mp11-12 SNP-based admixture plot is displayed between the dendrogram and heatmap. **E.** Rarefaction plot of homologs among all isolates or in selected groups, with and without singletons. **F.** Rarefaction plot of gene homologs for each population cluster. In both E and F, shaded regions surrounding the means represent one standard deviation above and below the mean with 1,000 randomized replications.

### *M. phaseolina* cluster XIII was specialized for virulence in strawberry and associated with recent outbreaks of strawberry crown rot disease

Significantly more isolates from strawberry (85 out of 106) were in cluster XIII than would be expected by random chance (χ^2^ *P* < 0.001). We hypothesized that cluster XIII isolates were specialized for virulence in strawberry and tested this with virulence assays of 41 isolates representing eight *M. phaseolina* clusters. Five of the six tested isolates from the strawberry-associated cluster XIII consistently caused disease in the susceptible strawberry cultivar, ‘Monterey’, as indicated by area under the disease progress curve (AUDPC) and measured plant and fruit biomass (Figure 3). No isolate from another cluster caused disease on strawberry plants, but almost all isolates could be recovered from asymptomatic plants at the end of the seven-week growth period (Figure 3D). This indicates that the potential for asymptomatic, endophytic growth on strawberry is common in *M. phaseolina*, but only cluster XIII is specialized for virulence on strawberry.

**Fig. 3.**
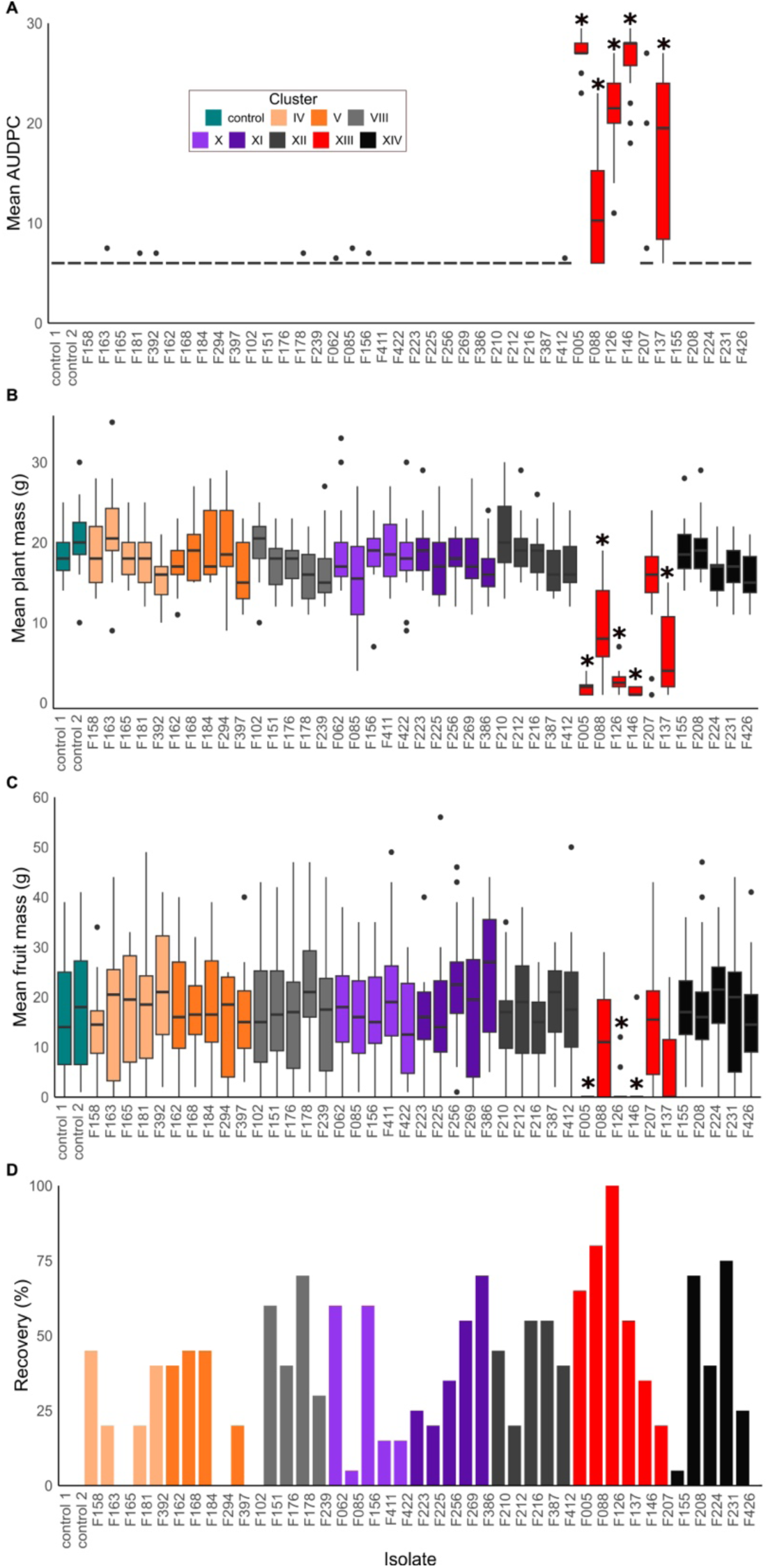
Virulence of *M*. *phaseolina* isolates in strawberry (cv. Monterey). Boxplots depict areas under the disease progress curve (**A**), above-ground plant fresh weight at the end of the experiment (**B**), and total fruit weight over the entire experiment (**C**). Plots depict the pooled results from two experiments, each with 10 biological replicates per treatment. Asterisks indicate a significant (*P* < 0.001) difference from the water control by a Mann-Whitney U test. **D.** Barplot depicting the percent recovery of *M. phaseolina* from all 20 plants per treatment.

The cluster XIII host-specialized strains appeared to have emerged recently and were responsible for most new reports of *M. phaseolina* causing disease in strawberry. Consistent with recent emergence, isolates in cluster XIII had low genetic diversity, as evidenced by their high fraction of gene conservation, and low anchor distance divergence (Figures 2D and 2F, and Supplementary Figure S4). Our dataset included only three isolates of XIII before it was first recovered from strawberry: chrysanthemum in 1934 from Missouri (F103), slash pine in 1983 from Paraguay (F287), and pepper in 2000 from Australia (F322). From 2001 onward, cluster XIII isolates were associated with the nearly simultaneous first reports of highly destructive disease in strawberry in Florida^50^, Spain^51^, California^52^, Mexico, and Australia^53,54^ (Supplementary Figure S5).

The emergence of this strain that is specialized for virulence in strawberry appears to be a major factor in the increased frequency and severity of strawberry crown rot disease caused by *M. phaseolina* worldwide. The emergence of this host-specialized strain also coincided with the withdrawal of methyl bromide as a highly effective soil fumigant. Many authors have postulated that *M. phaseolina* emerged in strawberries due to the discontinuation of methyl bromide as a highly effective soil fumigant, partly because epidemics mostly arose in fields that had adopted substitute fumigants^30,52,53^. This hypothesis is not universally supported by the data: side-by-side comparisons do not always show methyl bromide controlled *M. phaseolina* significantly more than replacement fumigants^30,53,55,56^. Furthermore, virulent isolates have been recovered from strawberry nurseries, where methyl bromide continues to be used^27^. Our data show that the emergence of a host-specialized strain was a crucial factor explaining new reports and the surging destructiveness of this pathogen in strawberry in multiple countries.

Biosafety restrictions limited us to testing isolates from the USA, so the strawberry-derived isolates from clusters XI, XII, and XIV were not tested. Some isolates in these three clusters were found to be virulent in previous studies and are associated with strawberry crown rot disease in Argentina^57^, Australia^58^, Chile^59^, Israel^60^, and Italy^61^. Thus, it is likely that other isolates of *M. phaseolina* could cause disease on strawberry, potentially under different environmental conditions and/or other cultivars than were tested in our study.

### Clusters IV, V, VI, VII, and VIII were associated with the soybean host

Most isolates from soybean or soil from soybean fields were assigned to five clusters: IV (7 soybean and soybean field isolates/9 total), V (34 isolates/41 total), VI (13 isolates/15 total), VII (31 isolates/38 total), and VIII (5 isolates/7 total; collectively 90/110). These distributions were significantly different from what would be expected under the null hypothesis of no association between the soybean host and isolate genotype (χ^2^ *P* < 0.001). The 110 soybean isolates were derived from the USA (56 isolates), Paraguay (42 isolates), Australia (5 isolates), and Denmark (1 isolate), indicating this trend appears to be a general phenomenon not limited to specific geographic areas. We hypothesize that the genotype × soybean host association results from increased virulence of these population clusters on soybean, as it was for strawberry.

The common refrain that *M. phaseolina* is a broad host range pathogen (reviewed in Pennerman, et al. ^9^) implies that the sub-specific genotype is irrelevant, as any isolate could cause disease in any host. Our data provide a compelling contradiction to this by demonstrating that a specific lineage, cluster XIII, is specialized for virulence on strawberry, and that specific clusters are associated with soybean. *Macrophomina phaseolina* is best characterized as both a poly-specialist, with multiple populations specialized for different hosts, and a generalist, with populations containing isolates derived from many hosts, no obvious specialization and a capacity for asymptomatic infection.

This is similar to what has been observed for *Botrytis cinerea*^62,63^ and *Magnaporthe oryzae*^2^, which provide useful paradigms for understanding host specialization in *M. phaseolina.* Lineages of *M. oryzae* are specialized for pathogenicity in specific grasses; when sampling for *M. oryzae* from natural blast infections, the host-specialized strain is almost always recovered^2,64^, similar to what we observed for strawberry and soybean. However, *M. oryzae* isolates retain virulence in grasses other than the one they are specialized in and can sometimes be recovered from other species^2,65^. We also observed a small percentage of strawberry- and soybean-derived isolates in other clusters and speculate that isolates in the strawberry- and soybean-associated clusters retain virulence in other hosts. Evaluating the host range of specific genotypes will be an important avenue of future research.

### Lack of genotypic association with geographic location suggests *Macrophomina* spp. were frequently trafficked between countries

Overall, SNP-based genotype clustering was not associated with the geographic origins of isolates (*P* = 0.616 after 10,000 permutations), as indicated by a Mantel test with the latitudes, longitudes and Q-values of 415 isolates with known sub-country regions. Similarly, analysis of molecular variance (AMOVA) identified more variation of Mp11-12 SNP alleles within countries (90.44%) than between countries (9.56%; *P* < 0.001) or between isolation sources (15.14%; *P* < 0.001). The average calculated fixation index (Hendrick’s corrected G_ST_, G’_ST_) among the genotype clusters (G’_ST_ = 0.439) was higher than when grouping isolates by country of origin (G’_ST_ = 0.155; Supplementary Figure S6), indicating that genetic diversity was lower when isolates were grouped by genotype than by geographic origin.

Similarly, most clusters were found in each of the three well-sampled continents. All SNP-based clusters were found in North America, from which most isolates were derived from USA (n = 220 isolates from the USA out of 230 total from North America). Clusters VI and VIII were exclusively found in the USA; these were the only clusters to be found only in one country or continent. The second-most sampled continent was South America (n = 86 isolates), which contained all clusters except VI, VIII and XIV. Australia was the third-most extensively sampled continent (n = 83), and it collectively contained all clusters except IV, VI, and VIII (Figure 4A). This indicates that isolates from each genotypic cluster are widely dispersed across continents rather than found specifically within certain regions. The weak influence of geography on population structure was visualized with minimum spanning trees generated from the Mp11-12 SNP data, which show isolates were more clearly defined by genotype than by host country (Figures 4B and 4C).

**Fig. 4.**
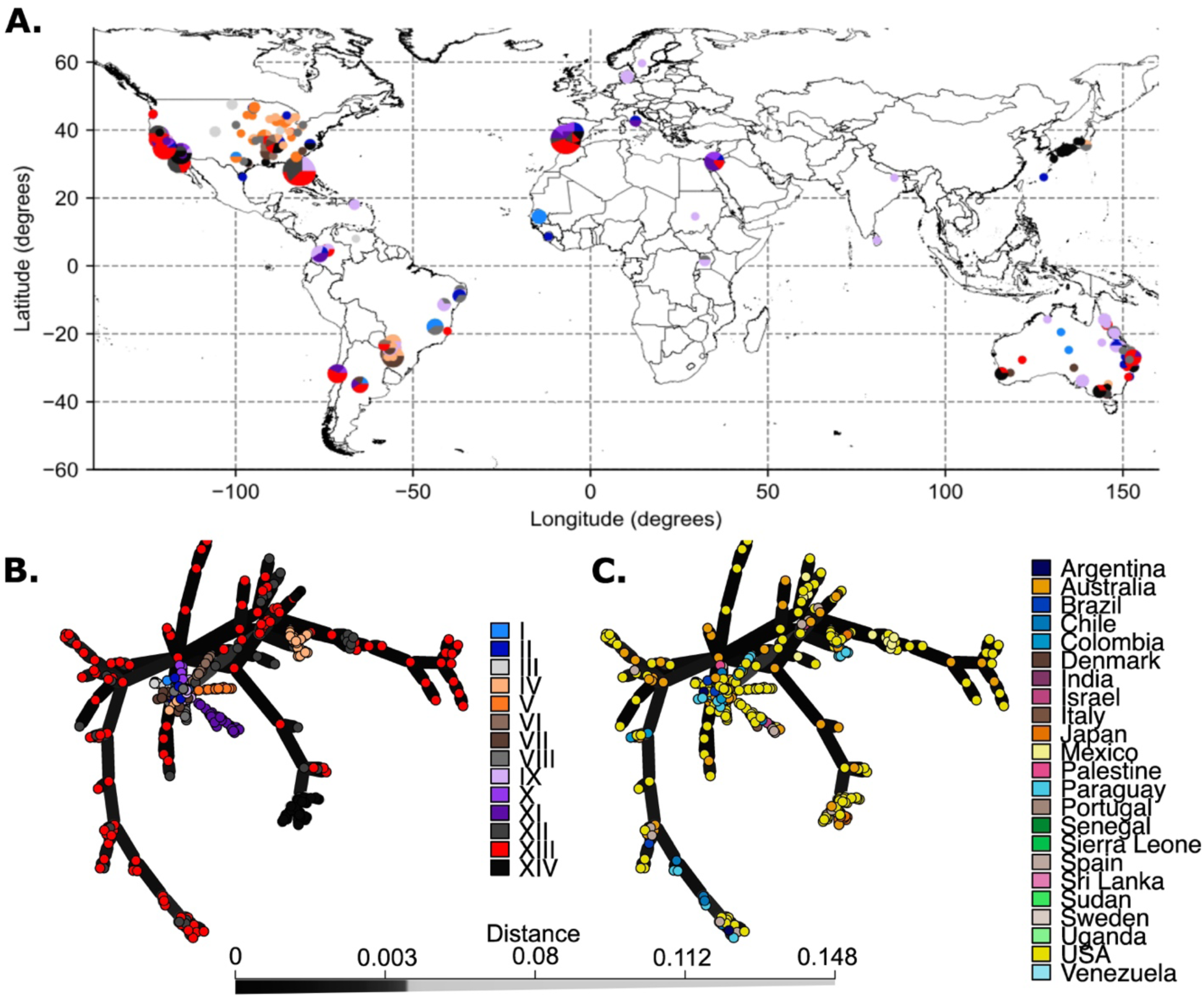
Geographic distribution of *Macrophomina* spp. genotype clusters. **A.** Isolates without a known state, province, or locality were assigned longitudes and latitudes at the midpoints of their respective sub-country region (Supplementary Table 1). Pie charts scaled to the number of isolates recovered are shown for locations that had multiple isolates. The color-code for the pie charts is the same as provided for panel B of this figure. Minimum spanning trees for isolates in this study colored by genotype cluster (**B**) or the country of origin (**C**).

The lack of geographic influence on population structure was surprising, given that many pathogens have population structures influenced by geography, even aerially dispersed fungi^3–5^. However, many Botryosphaeriaceae pathogens are known to asymptomatically colonize plant tissues^66^, and this niche has been considered a reason why populations of *Botryosphaeria dothidea* are not structured by geography^67^. The potential for asymptomatic growth on plant tissues or seeds by *M. phaseolina* is known for some hosts^28,33–36,38^, but is a relatively understudied phenomenon. In support of this hypothesis, our data showed that most *M. phaseolina* isolates could initiate asymptomatic infections in strawberry (Figure 3D). Similarly, we found that cluster XIII isolates of *M. phaseolina* could cause asymptomatic infections in a broad range of Fabaceae and Brassicaceae cover crops (Qin et al., in review). Asymptomatic infections on plants or seeds are likely to play a much larger role in *M. phaseolina*’s lifecycle than was previously appreciated and warrant greater attention in future research.

### Recombination appears to be ongoing in *Macrophomina* spp., including between *M. phaseolina* and *M. tecta* and, and could be the result of meiosis

Evidence of apparently recent and on-going recombination was revealed by fractional ancestries (quantified by Q-values) and LD decay. Most isolates (361 out of 463) had high (>95%) fractional ancestries, meaning that most of their SNP genotype calls corresponded to their assigned cluster (Supplementary Table S1). However, there were also 57 isolates that were admixed, with less than 75% of their fractional ancestry attributed to their assigned cluster, and 30 isolates that were highly admixed with less than 50% of their fractional ancestry represented by their assigned cluster. In LD decay plots, the *r^2^* values declined to around 0.15, after which they tended to remain level, similar to what has been shown for populations with on-going recombination (Supplementary Figure S7)^3^. There was evidence for hybridization between *M. tecta* (cluster II) and *M. phaseolina*, as several cluster II isolates had high ancestry fractions for *M. phaseolina* clusters (Figure 1A; Table 1). Evidence for hybridization between cluster I (*M. pseudophaseolina* and *M. euphorbiicola*) and *M. phaseolina* clusters was less apparent, with Q-values for the assigned cluster among cluster I isolates being at least 90.4% for the Mp11-12 reference. This suggests that recombination could be occurring between *M. tecta* and *M. phaseolina*, but may not be occurring between species in cluster I and *M. phaseolina*.

**Table 1.**
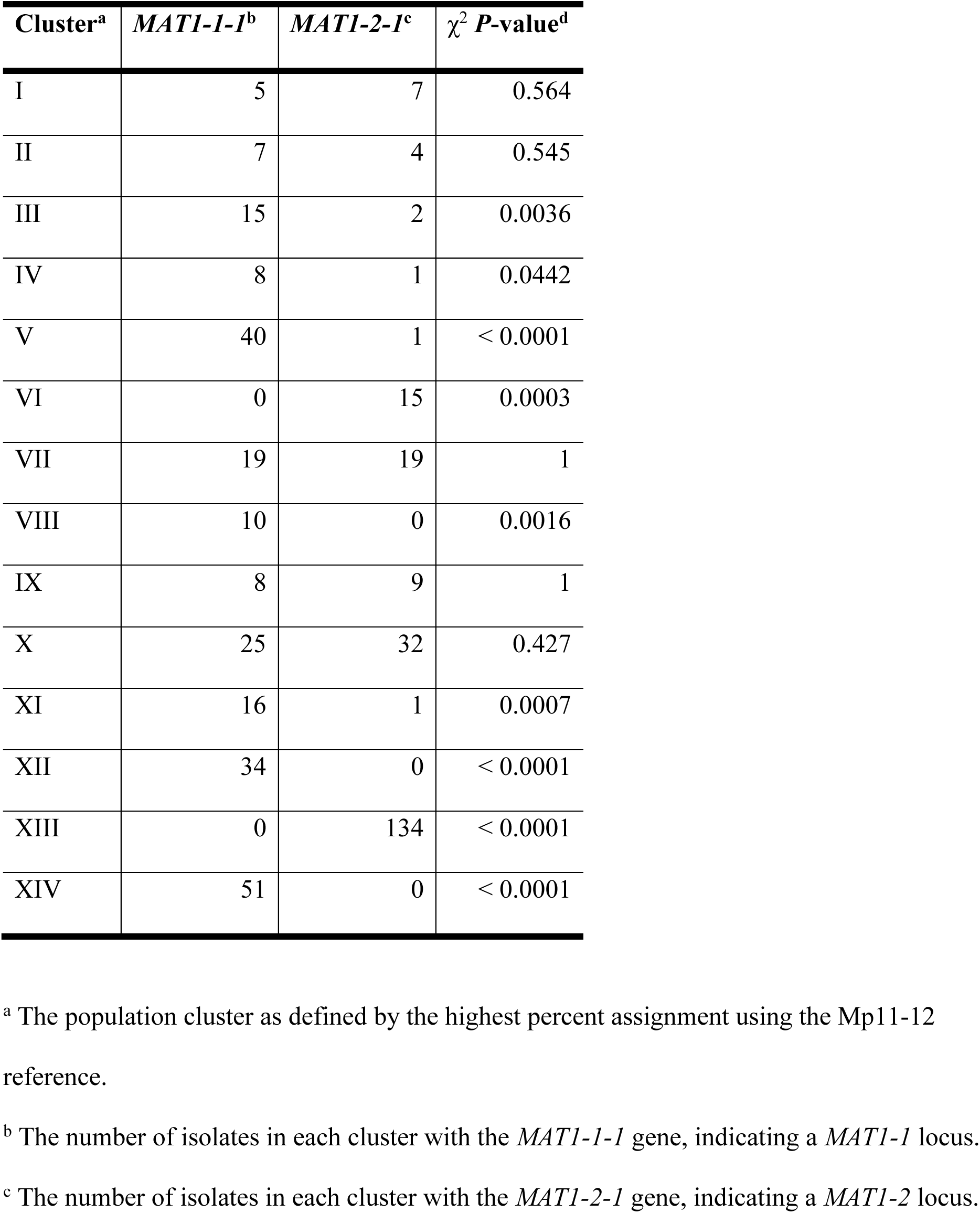
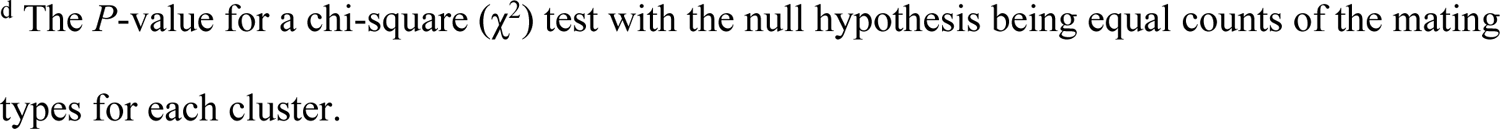
Mating type ratios of *Macrophomina* spp. genotype clusters.

A heterothallic mode of sexual reproduction for *M*. *phaseolina* was previously reported by Nagel et al.^44^. This was confirmed here for *M. phaseolina* and newly determined for *M. pseudophaseolina*, *M. euphorbiicola*, and *M. tecta*. Across all isolates there were nearly equal numbers of each mating type (mating type 1 = 238, mating type 2 = 225), but mating type distribution was significantly different from the expected 1:1 ratio for nine clusters (χ^2^ *P* = 0.0442; Table 1). Five clusters had equal mating type ratios (χ^2^ *P* = 0.427; Table 1). While admixed isolates overall had partial memberships in all clusters (Supplementary Table S2), four of these clusters, II, III, VII, and IX, also had relatively low average Q-values corresponding to the main genotype for all references (Supplementary Table S4). This suggests that isolates in these populations were more prone to “outcrossing” with isolates from other clusters. These data indicate that the genetic potential for meiosis exists in *Macrophomina* spp. and is likely to contribute to recombination in these species.

### Genomic evidence supports the delineation of currently described *Macrophomina* spp., which can be differentiated by new species-specific primers

Reference-independent, whole genome comparisons of anchor nucleotide distances (ANDI) supported the circumscription of *M. phaseolina*, *M. pseudophaseolina*, *M. euphorbiicola*, and *M. tecta*. The minimum ANDI for any two isolates of different species (= 0.0113) was greater than the maximum ANDI for any two isolates of the same species (= 0.009; Supplementary Figure S4). There was just one isolate (F289) that did not fall into any of these species, as its pairwise ANDI values were all ≥ 0.0113; above the threshold observed for within-species comparisons. A multi-locus tree based on a concatenation of four commonly-used phylogenetic markers showed that F289 was most similar to the type strain of *M. vaccinii* (Supplementary Figures S8 and S9). Morphological comparisons between F289 and selected *M. phaseolina* strains showed pycnidia or conidia sizes overlapped with those reported for both *M. phaseolina* and *M. vaccinii*^20^ (data not shown), indicating that these characteristics could not clearly differentiate these species. We considered F289 to be a putative *M. vaccinii* isolate but could not confirm this with genomic comparison to known strains of this species.

A series of species-specific qPCR assays were designed and validated to enable rapid identification of the species analyzed in this study. The qPCR assays were first validated *in silico* to target an identified locus that was present in each tested species (*phaseolina*, *pseudophaseolina*, *euphorbiicola*, and *tecta*) but absent in off-target organisms in Botryosphaeriaceae and the nr/nt database of Genbank. Each target was tested *in vitro* on a panel of 97 isolates and found to generate the predicted results for all assays, with one exception: the *M. tecta* assay also amplified F289, the one isolate that could not be classified to species.

Additionally, the *M. phaseolina* assay did not amplify an individual *M. phaseolina* isolate that was highly admixed with *M. tecta*, as expected (Supplementary Tables S1, S2 and S3). These results highlight the limitations of species boundaries in the genus *Macrophomina*, which may be blurred by inter-specific hybridization.

We also designed a conventional PCR assay that generates a 507 bp amplicon, here called the “mspp locus”, that can be sequenced and aligned to known references. This locus is within a predicted fatty acid synthase beta subunit, annotated as gene “M11_12_v1_07558” in Burkhardt, et al. ^68^. The mspp PCR assay was validated with representatives of each *Macrophomina* spp.; the amplicons were sequenced and yielded the expected result (Supplementary Figure S10). Alignment of sequences at the mspp locus can differentiate all *Macrophomina* species and the population cluster XIII with a high degree of accuracy (Table 2; Figure 5). The mspp locus provided a similar amount of phylogenetic information as the standard approach of sequencing and concatenating alignments of ITS, *TEF1-α*, *actin*, and *beta-tubulin* sequences, and it is more informative than any of these standard reference genes on their own (Figure 5; Supplementary Figure S9). Notably, the ITS sequence was not sufficient to delineate *Macrophomina* spp., as multiple species are represented by single ITS haplotypes (Supplementary Figure S8).

**Fig. 5.**
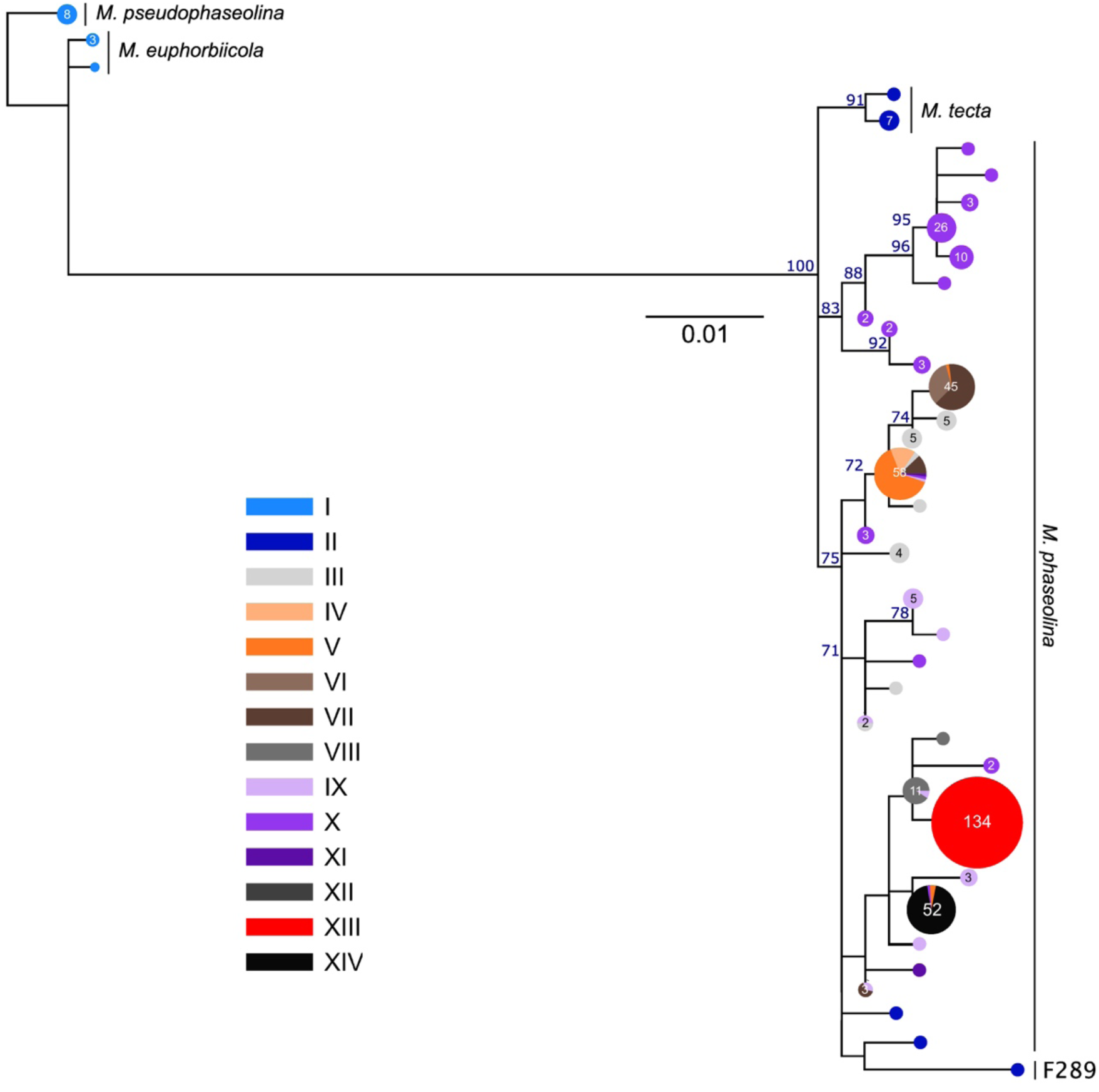
Phylogenetic tree built from sequences at the ∼500 bp locus targeted by the “mspp” PCR primers. The phylogenetic tree was calculated by RAxML-NG version 1.2.1 with the GTR+G model of evolution and 1,000 bootstrap replicates. Only bootstrap values greater than 70 are shown. Species *M*. *euphorbiicola*, *M*. *pseduophaseolina*, *M*. *tecta*, and *M*. *phaseolina* are distinguishable and indicated by text annotations. All isolates from the strawberry-associated cluster XIII had identical sequences that were distinct from other *Macrophomina* spp. isolates. Used sequences are in Supplementary Information S3.

**Table 2.**
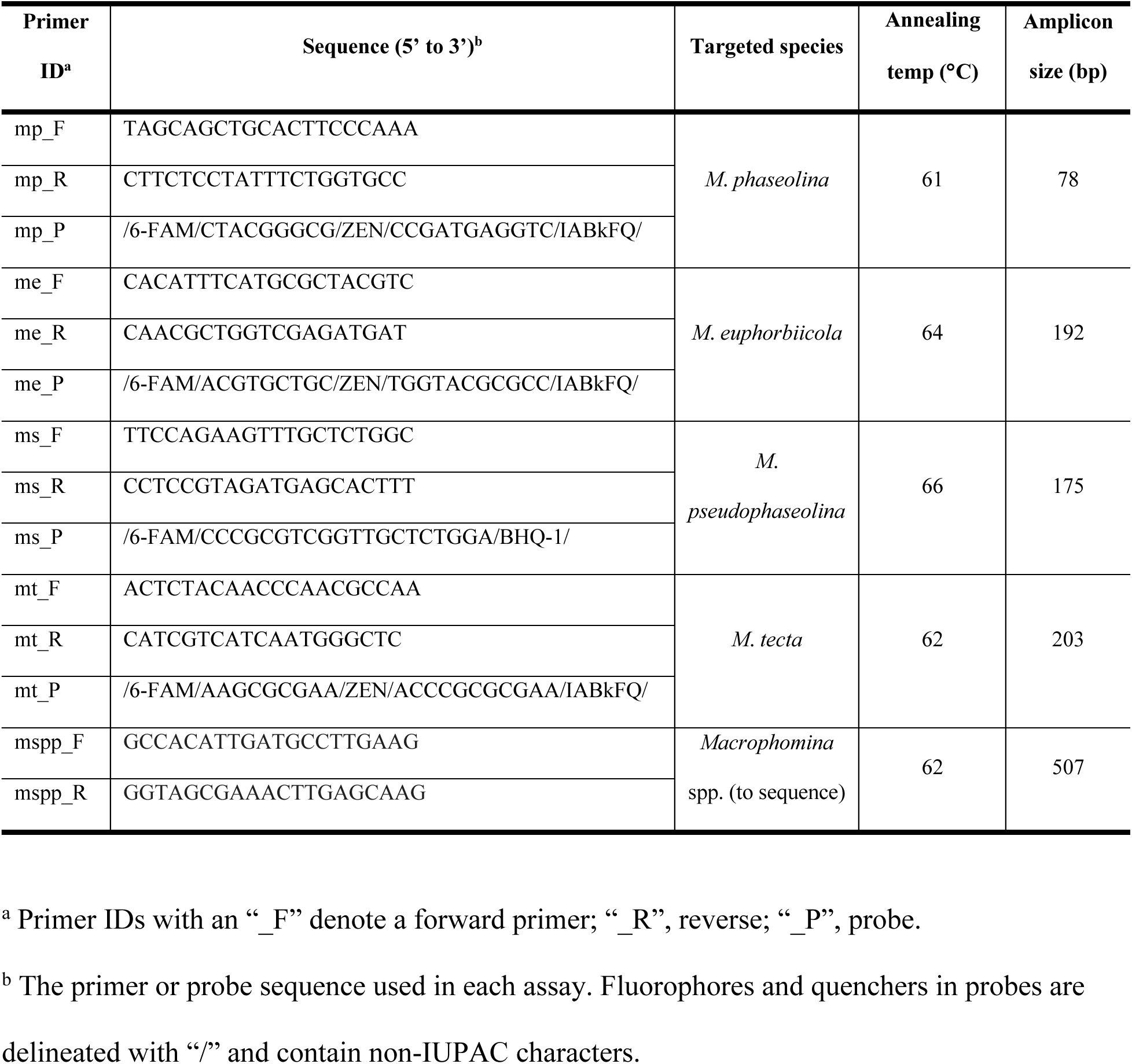
Primers and probes designed for detection and differentiation of *Macrophomina* spp.

The currently described *Macrophomina* species were supported by reference-free whole genome comparisons (Supplementary Figure S4) and phylogenies of core genes (Supplementary Figure S9). However, admixture between species indicate that the boundaries in *Macrophomina* spp. are fluid and could complicate future efforts at specific detection. This led to challenges designing qPCR assays with 100% analytical specificity, as an admixed isolate with *M. phaseolina* and *M. tecta* ancestry did not amplify with either assay. However, our species-specific assays performed well overall and were the first of their kind for *M. pseudophaseolina*, *euphorbiicola*, and *tecta*. The *M. phaseolina* assay provides a slight improvement on the existing qPCR assay^69^ by including a TaqMan probe in the design, which enables multiplexing with other targets. The existing *M. phaseolina* assay by Babu, et al. ^69^ was *in silico* predicted to correctly amplify all *M. phaseolina* isolates and not other *Macrophomina* species. Our new *M. phaseolina* assay also has an intentionally broader target than the Burkhardt, et al. ^70^ assay, which is only predicted to amplify all isolates in cluster XIII except F219.

## Methods

### Global collection and sequencing of *Macrophomina* spp. isolates

Effort was made to collect isolates from diverse hosts, geographic locations, and times. Isolates were drawn from public culture collections (e.g. the Queensland Herbarium, Westerdijk Institute, NARO Genebank, and the National Reference Repository Library), private collections of individual investigators, and recently published works^27,71^. A standardized naming system was applied to 463 isolates received, ranging from F001 to F518. Metadata for each isolate were largely provided by the collectors and are listed in Supplementary Table S1. Longitudes and latitudes were assigned to isolates based on the midpoint of known states, provinces, or localities.

### DNA extraction and sequencing

All isolates were single-hyphal-tipped to ensure genetic uniformity. The tissue used for DNA extraction was grown on potato dextrose agar, potato dextrose broth, or 10% V8 juice broth, and was sometimes lyophilized. DNA extractions were performed with the OmniPrep for Fungus kit (G-Biosciences, USA) per manufacturer’s instructions, with an added step of treatment by Invitrogen RNase A/T1 Cocktail (Fisher Scientific, Massachusetts, USA) at the final step and incubating for 60 minutes at 45°C followed by 30 minutes at room temperature. Most DNA extracts were then further purified by AMPure XP beads (Beckman Coulter, California, USA) to improve the quality of DNA extracts before library preparation. Libraries were prepared with the DNA *KAPA Hyper Prep Kit* (Kapa Biosystems-Roche, Basel, Switzerland) and sequenced on an Illumina NovaSeq6000 with 150 bp paired-end reads. For isolates from Brazil, DNA was extracted by the method described in Kjærbølling, et al. ^72^ and sequenced on an Illumina NextSeq2000 at the Max Planck Genome Center in Cologne, Germany. Additional reads were retrieved from NCBI GenBank for 28 isolates; accession numbers are available in Supplementary Table S1. Estimated genomic sequencing coverage was typically between 72× and 108×, while a small number of isolates had coverage between 26× and 64× (Supplementary Table S1).

### Sequence read processing and mapping

Unless specified, libraries, modules and codes were used at their default parameters. Sequencing reads were preprocessed by removing contaminants, unique molecular identifiers, adaptors, ambiguous and low-quality nucleotides using HTStream version 1.3.3 (https://github.com/s4hts/HTStream). BBMap version 35.85 (https://sourceforge.net/projects/bbmap/) was used to limit the total number of bases to a maximum of 5.5 Gb (approximately 100× coverage) with a minimum read length of 100 bases for each isolate. Cleaned reads were mapped to the assembled genomes of *M*. *phaseolina* strains Mp11-12 (F001), AL-1 (F003) and mp117 (F509)^45,68,70^ using the Burrows-Wheeler Aligner version 0.7.17^73^ and alignments were subsequently processed with SAMtools version 1.16.1^74^. BEDtools version 2.30.0^75^ converted the resulting BAM files into BED files. SAMtools flagstat evaluated mapping efficiency. The BED files of mapped read coverages were used to determine if the strawberry *M. phaseolina* detection locus^70^ and the general *M. phaseolina* detection locus^69^ were present among the isolates by counting at least five sequencing reads mapped to a total of at least 95% of the length of the reference locus.

### Genome assembly and annotation

Cleaned reads were *de novo* assembled into genomes with SPAdes version 3.15.5^76^, and genome assembly quality was evaluated using QUAST version 5.2.0^77^ and BUSCO version 5.4.5^78^ with the dothiodeomycetes_odb10 database (version 2019-11-20), which contained 3,786 benchmarking universal single-copy homologs. SPAdes assemblies were processed through the NCBI Foreign Contamination Screen to remove sequencing adapters and prokaryotic contaminants before submission to NCBI WGS^79^.

Protein-coding genes were annotated from these assemblies with GeneMark-ES version 4.71^80^ with the fungus parameter. Proteins with fewer than 100 amino acids were excluded from analyses. These putative proteins were consolidated into a pan-proteome with CD-Hit version 4.8.1^81^ with the reciprocal sequence identity threshold and length difference thresholds at 0.8. “Super-core” homologs were represented by every isolate included in the current study. “Core” homologs included super-core homologs and those that were present in at least 95% of isolates. “Accessory” homologs were present in at least 2 isolates and up to 95% of isolates. “Singleton” sequences were only identified in one isolate.

### Variant calling and population genomic analyses

Biallelic SNPs with a minimum coverage of five reads for each isolate, a minimum base quality of 20 for each isolate, and a minimum allele frequency of 5% overall among all *Macrophomina* spp. isolates were identified with Freebayes version 1.3.6^82^. SNP loci missing from more than 10% of used *Macrophomina* spp. isolates were excluded. Admixture was analyzed with the unpruned loci with sNMF version 1.2^83^ with K ranging 2 to 20. A final K of 14 was chosen in accordance with the elbow method for clustering algorithms. SNP intra-scaffold LD correlations were calculated with VCFTools^84^ and the unpruned loci. Fixation indices were computed with R package vcfR^85^. The R package poppr was used to generate minimum spanning trees and to conduct AMOVAs^86^. Sequences corresponding to the *MAT1-1-1* and *MAT1-2-1* genes were extracted from full mating type loci published by Nagel, et al. ^44^ and then BLASTn searches of these queries against each reference assembly were used to determine which gene was present in each isolate.

Taxonomic assignments were verified by retrieval of reference nucleic sequences of ITS (reference NCBI accessions KF951717.1, KF951791.1, OR052068.1, MK687450.1 and NR_175162.1), *TEF1-α* (KF952088.1, KF952153.1, OR030468.1, MK687426.1 and MW592271.1), *actin* (KF951857.1, KF951918.1, MH712508.1, MK687442.1 and MW592058.1), *beta-tubulin* (KF952178.1, KF952233.1, MF457658.1, MK687434.1 and MW592300.1) genes or genomic regions. These reference sequences represented phylogenetic markers from *M*. *phaseolina* CPC 21420, *M*. *pseudophaseolina* CPC 21417, *M*. *tecta* BRIP:70781, *M*. *vaccinii* CGMCC3.19503, and *M*. *euphorbiicola* strains CMM4134, McBr87 and MFLU 23-0004. The reference sequences were used in BLAST searches^87^ against our assembled genomes to identify the corresponding sequence in each isolate. The ITS, *TEF1-α*, *actin*, and *beta-tubulin* sequences were aligned by MUSCLE^88^ and RAxML-NG version 1.2.1^89^ was used to generate phylogenetic trees with the GTR+G evolution model and 1,000 bootstrap replicates. Phylograms were visualized with the R package: ggtree^90^. The software andi^91^ was also used to estimate isolate relatedness with the assembled genomes. Reticulate phylogenetic trees were generated using SplitsTree version 4.19.1^92^ (NeighborNet function). Other figures were made with Inkscape 1.3, Python 3.11.4 libraries matplotlib and sklearn, and R 2023.03.1. Heatmaps were generated using Ward’s method parameter in the Python Seaborn library and using data that was one-hot encoded. GIS shapefiles were retrieved from the IPUMS database^93^.

### Development of species-specific qPCR assays and a PCR protocol for a phylogenetically-informative locus

Phylogenetically-informative sequences commonly shared among all isolates were identified using the BEDTools suite^75^ to compare BED files of read coverages to the Mp11-12 genomic reference. Species-specific target candidates for qPCR assays were identified by searching the CD-HIT determined homologs for those that were specific to the target *Macrophomina* spp. Putative species-specific homologs were further verified by BLAST searches of the locus against all genome assemblies and NCBI nr/nt and wgs sequence databases.

Species-specific Taqman qPCR assays were designed for *M. phaseolina*, *M. pseudophaseolina*, *M. euphorbiicola*, and *M. tecta*. All assays used a 20 µL reaction mixture containing 1× PerfeCTa qPCR ToughMix (Quantabio, Massachusetts, USA), 0.25 µM of forward and reverse primers (Table 2), 0.15 µM of Taqman probe (Table 2), and approximately 15 ng of DNA. DNA for these reactions was isolated by the OmniPrep kit as described above or by the Quick-DNA Miniprep kit (Zymo Research, California, USA). Reactions were run in a LightCycler II (Roche, USA) with an initial denaturation step of 95°C for 2 min followed by 40 cycles of 15 sec at 95°C and 30 sec at the annealing temperatures provided in Table 2. These qPCR assays were tested on 97 isolates that included at least two representatives of each species and at least five representatives of each *M. phaseolina* population cluster (Supplementary Table S3). All qPCR assays were predicted to yield exactly zero or one amplicon depending on the isolate (Supplementary Table S1).

An additional protocol was developed for PCR amplification of a single, phylogenetically-informative locus present in all *Macrophomina* spp. that can be sequenced to determine species assignment. This assay was run with a 50 µl reaction mixture containing 0.5 µM of each primer (Table 2), 1× Promega Go Taq Master Mix (#M7123, Promega, Wisconsin, USA), and 2 µL of DNA isolated using the Quick-DNA Fecal-Soil Microbe kit or the Quick-DNA Miniprep kit (Zymo Research, Irvine, CA). Reactions were run in a BioRad C1000 Touch (Bio-Rad Laboratories, California, USA) with an initial denaturation step of 95°C for 5 min followed by 40 cycles of 95°C for 30 sec, 62°C for 30 sec, and 72°C for 1 min, then a final 72°C extension for 5 min. PCR products were purified using the GeneJet PCR Purification Kit (Thermo Fisher Scientific, New Hampshire, USA) and the 507 bp bands were visualized with GelRed Nucleic Acid Stain (Biotum, California, USA) on a 1.5% agarose gel for 40 minutes at 110V. Purified amplicons were sequenced with forward and reverse primers by Eton Biosciences (California, USA).

### Virulence assays in strawberry

Due to biosafety permit restrictions, we could only test virulence in strawberry for isolates from the USA in the greenhouse environment. Therefore, USA-derived isolates were selected at random from most *M. phaseolina* phylogroups: IV, V, VIII, X, XI, XII, XIII, and XIV. Isolates were grown in the dark for one week on potato dextrose agar amended with 50 mg/mL kanamycin, 100 mg/mL streptomycin and 12.5 mg/mL chlortetracycline. One 5-mm diameter plug of each culture was transferred to 200 mL of sterile 10% V8 broth and incubated at 25°C and 200 rpm in the dark for three days. The liquid cultures were then blended in a Waring blender for seven seconds and allowed to settle for five minutes in a 50 mL conical tube before the volumes of settled mycelia were quantified. Suspensions were diluted to 8% mycelia by volume with sterile deionized water.

Bare root, frozen strawberry crowns (cultivar Monterey) from a low-elevation nursery were planted in potting soil (Sunshine Mix #1, Nutrien Ag Solutions, Colorado, USA) and grown at 25°C/20°C day/night temperatures in a greenhouse with no supplemental lighting for one week to break dormancy. After dormancy was broken, the roots were rinsed with tap water and submerged in fungal inoculum for ten minutes. The negative control was submerged in 10% V8 broth for ten minutes. Plants were then repotted in 10.8 cm x 10.8 cm x 12.4 cm pots and watered with the respective leftover inoculum. Inoculated plants were then grown at approximately 28°C/22°C in a greenhouse with day lengths between 14 and 14.6 hours in Salinas, California. For the course of the experiments, plants were watered as needed and fertilized weekly using Peter’s Professional 20-20-20 fertilizer (ICL Fertilizers, Missouri, USA). Ten biological replicates for each fungal isolate were established per experiment, and the experiment was conducted twice.

Ordinal disease ratings were conducted weekly for seven weeks on a 1-5 scale, where 1 = asymptomatic, 2 = stunting, 3 = wilting of less than 50% of the foliage, 4 = wilting of more than 50% of foliage, and 5 = plant death. All peduncles bearing fruit and flowers were removed and weighed at five- and seven-weeks post-inoculation. After removing peduncles, fruit, and flowers for separate measurements at seven weeks post-inoculation, whole plants were removed from the soil, roots were trimmed flush with the crown, and above-ground fresh weight was recorded.

From each plant, two cross-sections of the crown were excised and their cortical tissues removed (as per Henry et al.^94^). These crown stele tissues were surface sterilized in 1% sodium hypochlorite solution for two minutes, then plated on potato dextrose agar amended with kanamycin, streptomycin, and tetracycline as described above. Plates were incubated at 30°C for 5-7 days, and the presence of *M*. *phaseolina* determined by the morphology of fungal growth emanating from these tissues. Data were analyzed by checking for normality with the Shapiro-Wilk and Levene’s tests. The Wilcoxon test identified statistical significance at α = 0.05.

## Author contributions

P.M.H. conceptualized the study. P.M.H., P.G., M.A., C.B., A.G., J.N., M.C., V.O., E.H.S, D.P., G.H.G., A.M., H.D.L.-N., N.V., L.K., J.P.B., A.R.M., T.S., N.A.P., J.B., K.I., G.C., S.K., D.M., S.M., and M.G. obtained and curated isolates for sequencing. P.M.H., G.H.G., J.P.B., A.R.M., D.P., E.H.S., and M.G. conducted whole genome sequencing. K.K.P, P.G., C.J.D and P.M.H performed data curation. K.K.P., P.G., C.J.D, G.R., J.H.J., J.L.-H., J.R., G.O.S., and P.M.H. conducted lab work including DNA extractions and pathogenicity study designs and/or executions. K.K.P. and P.M.H. analyzed data and wrote the manuscript. All authors edited and approved the manuscript.

## Data availability

Raw sequencing reads were deposited in the NCBI SRA database with BioProject accessions PRJNA953043. Individual isolate metadata and SRA accessions are included as part of Supplementary Table S1. Code associated with the analyses reported here are available at the Zenodo repository https://zenodo.org/records/13367157 with DOI: 10.5281/zenodo.13367156.

## Acknowledgements

We thank our funding sources, including the United States Department of Agriculture (USDA) Agricultural Research Service (ARS), the National Institute of Food and Agriculture Specialty Crops Research Initiative (#2017-51181-26833 and #2022-51181-38328) and the Technical University of Munich - TUM Institute for Advanced Study, funded by the German Excellence Initiative for a Hans Fischer fellowship awarded to G.H.G. This research was supported in part by an appointment to the ARS Research Participation Program administered by the Oak Ridge Institute for Science and Education (ORISE) through an interagency agreement between the U.S. Department of Energy (DOE) and the USDA. ORISE is managed by ORAU under DOE contract number DE-SC0014664. All opinions expressed in this paper are the authors’ and do not necessarily reflect the policies and views of USDA, DOE, or ORAU/ORISE.

## Supplementary Information Legends

**Supplementary Information S1.** Multiple sequence alignment of the ITS regions of *Macrophomina* spp. isolates and reference genomes. Sequences were de-duplicated, and headers contain the IDs of represented isolate(s) and/or reference(s).

**Supplementary Information S2.** Multiple sequence alignment of the concatenated ITS, *actin*, *beta-tubulin*, and *TEF1-α* sequences of *Macrophomina* spp. isolates and reference genomes. Sequences were de-duplicated, and headers contain the IDs of represented isolate(s) and/or reference(s).

**Supplementary Information S3.** Multiple sequence alignment of the mspp region of *Macrophomina* spp. isolates. Sequences were de-duplicated, and headers contain the IDs of represented isolate(s).

