## Supplementary Figures for "Population genomics of *Macrophomina* spp. reveals cryptic host specialization and evidence for meiotic recombination"

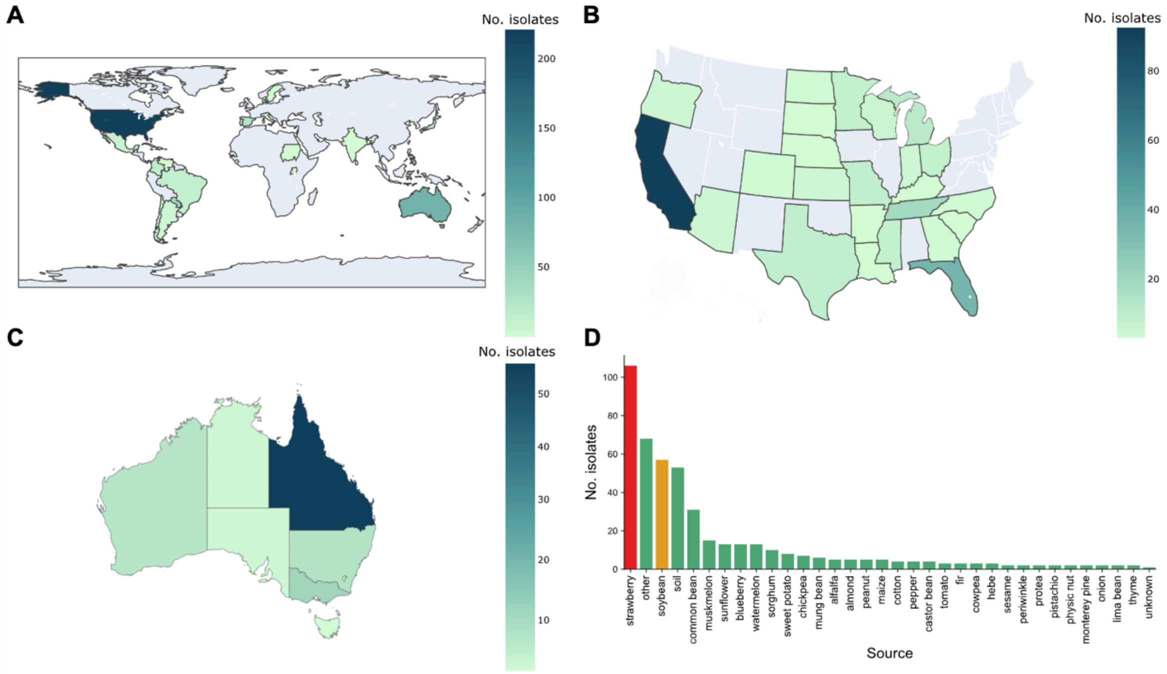


**Supplementary Fig. S1. Sources of *Macrophomina* spp. isolates. A**, **B**, **C** Geographic and **D** host origins of *Macrophomina* spp. isolates analyzed in the current study. Though **A** global collections, the plurality of isolates was collected in **B** the continental United States of America or **C** Australia, and **D** from strawberry or soybean plants. Choropleth colors correlate to the number of used isolates from a geographic region. Bar graph colors red and orange signify strawberry and soybean hosts.

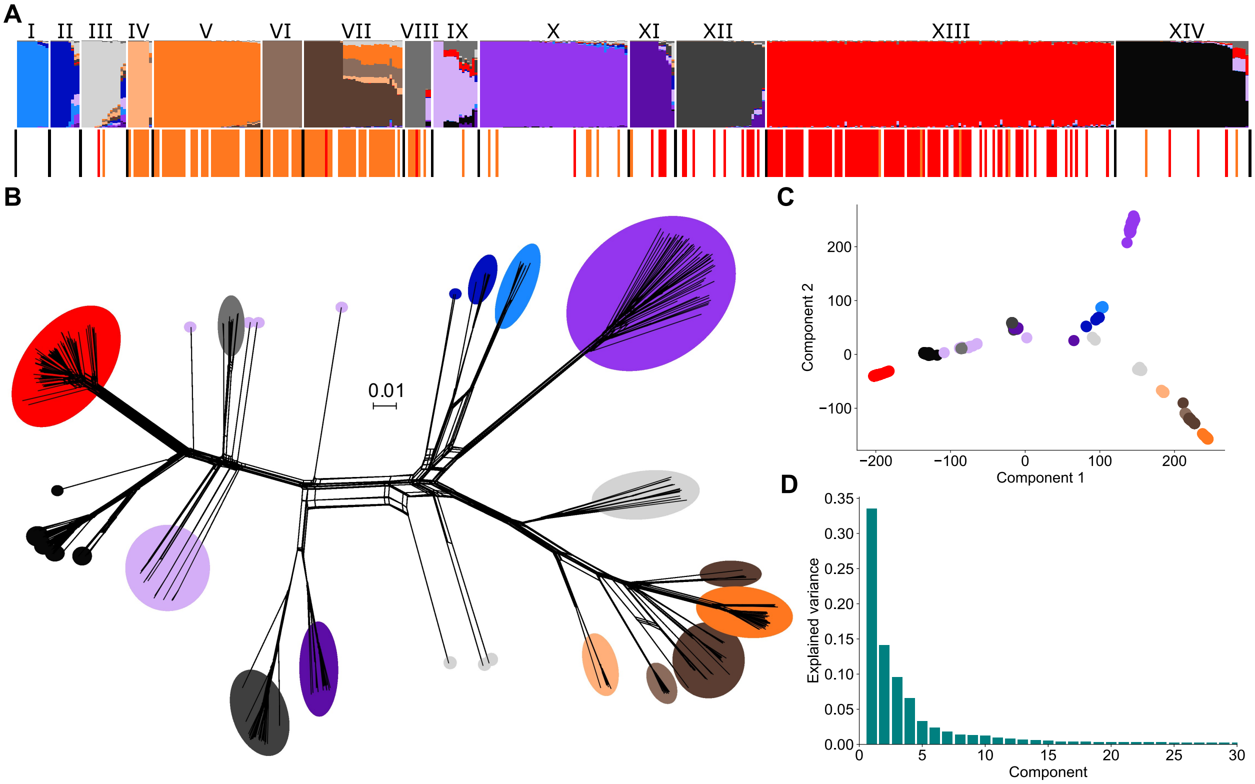


**Supplementary Fig. S2. Grouping of *Macrophomina* spp. isolates with AL-1-based SNPs.** The **A** admixture plot was generated with sNMF at K=14. From left to right, the clusters are labeled with Roman numerals corresponding to those used in the main text, with Mp11-12. The **B** phylogenetic network and **C** PCA plots use the same colors as the admixture plot to indicate the major cluster assignment. The **D** Scree plot shows the eigenvalues for the first 30 principal components.

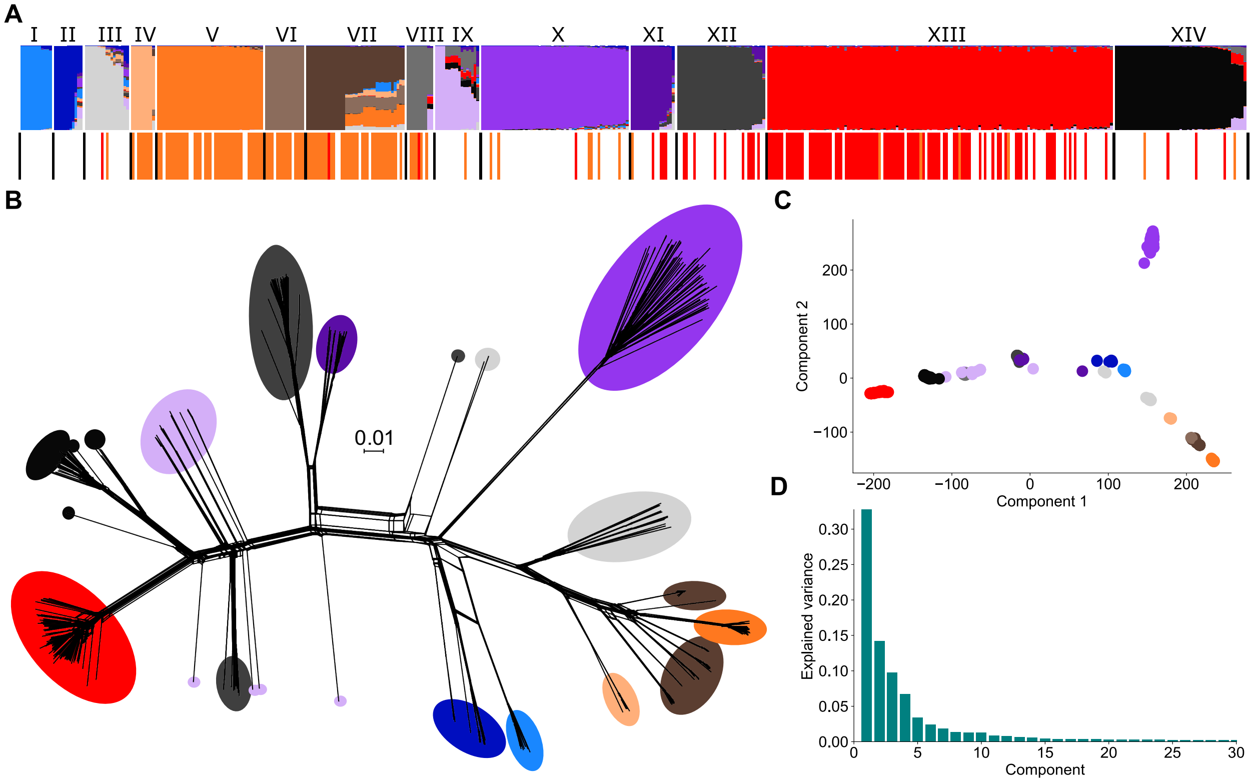


**Supplementary Fig. S3. Grouping of *Macrophomina* spp. isolates with mp117-based SNPs.** The **A** admixture plot was generated with sNMF at K=14. From left to right, the clusters are labeled with Roman numerals corresponding to those used in the main text, with Mp11-12. The **B** phylogenetic network and **C** PCA plots use the same colors as the admixture plot to indicate the major cluster assignment. The **D** Scree plot shows the eigenvalues for the first 30 principal components.

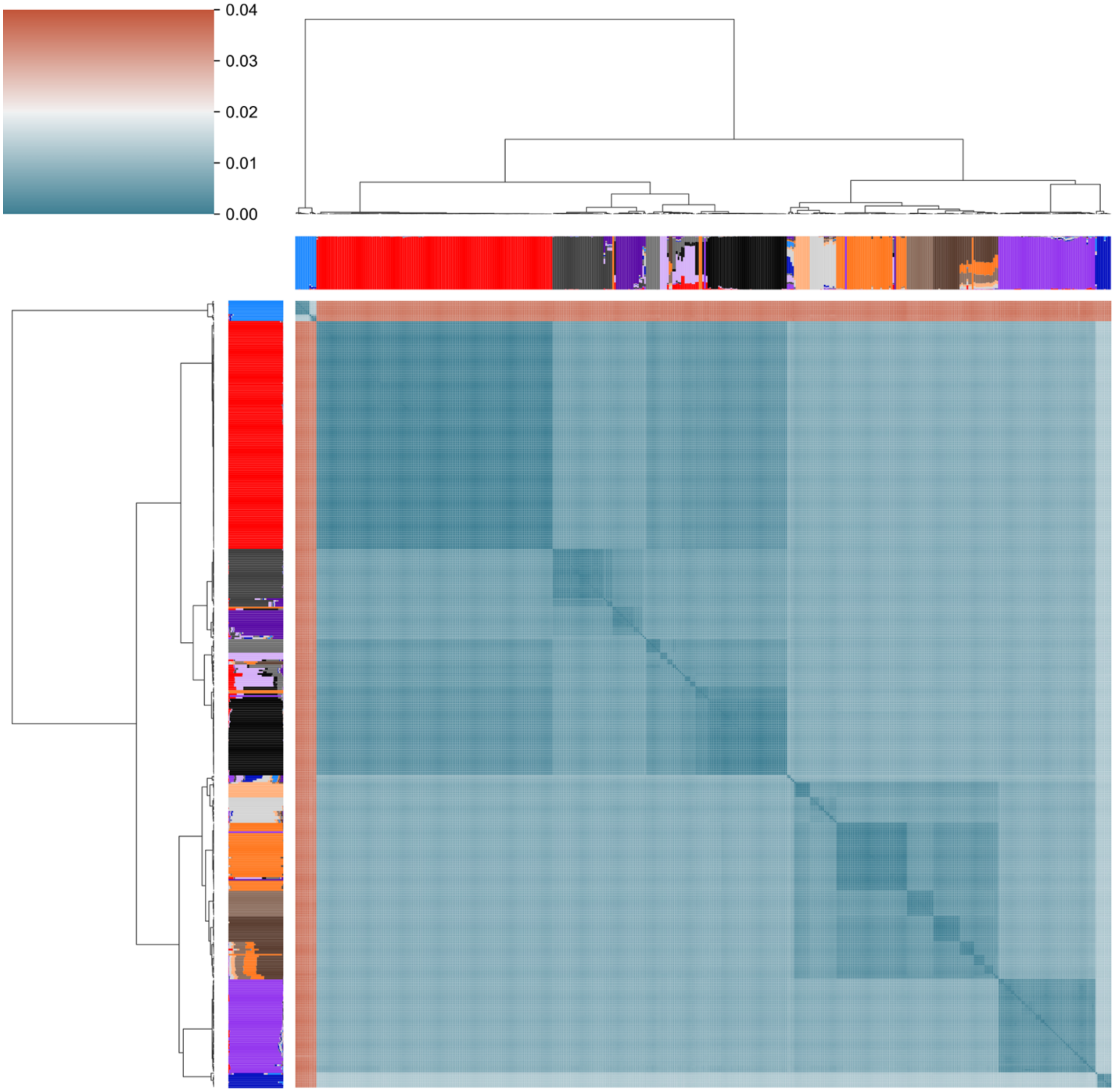


**Supplementary Fig. S4. *Macrophomina* spp. isolate relatedness according to pairwise anchor distances (ANDI).** Color value and intensity within the heatmap correspond to distance between a pair of isolates. Higher values indicate more dissimilarity. The corresponding Mp11-12 SNP-based admixture plot is overlaid on top utilizing the usual cluster-specific colors: I- light blue, II- dark blue, III- lightest grey, IV- light orange, V- orange, VI- light brown, VII- brown, VIII- grey, IX- lightest purple, X- purple, XI- darkest purple, XII- darkest grey, XIII- red, and XIV- black.

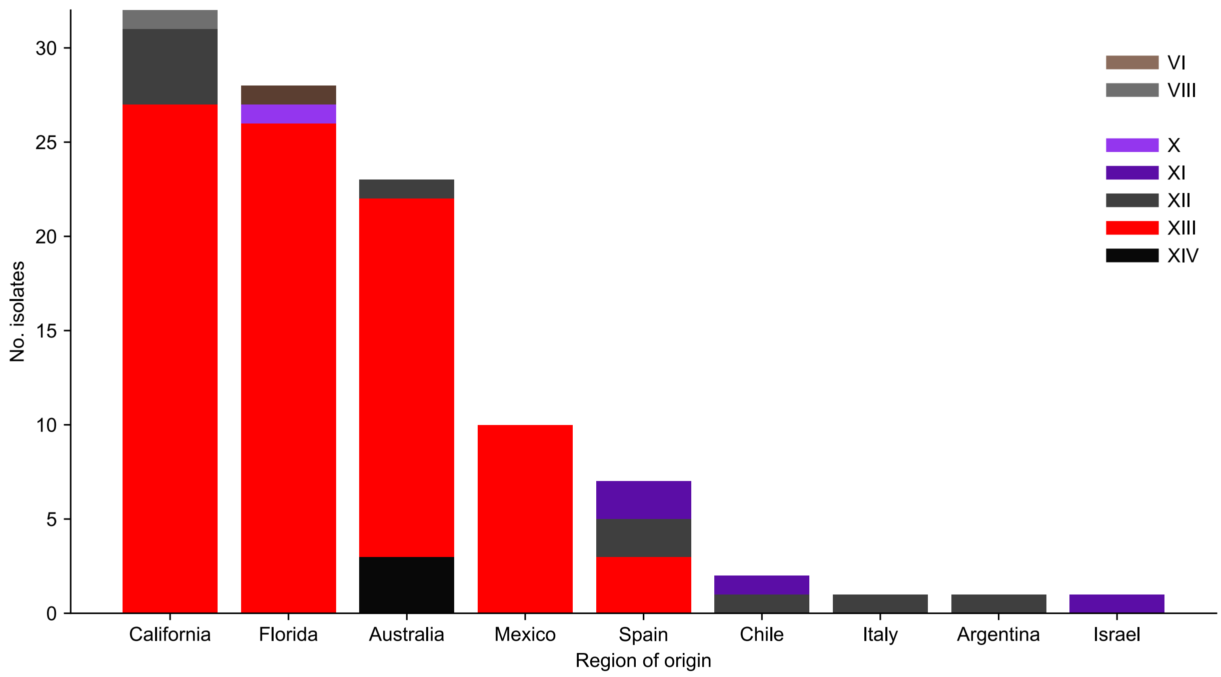


**Supplementary Fig. S5. Cluster assignments of isolates from strawberry by host country or USA state.** Most isolates from strawberry were assigned to cluster XIII.

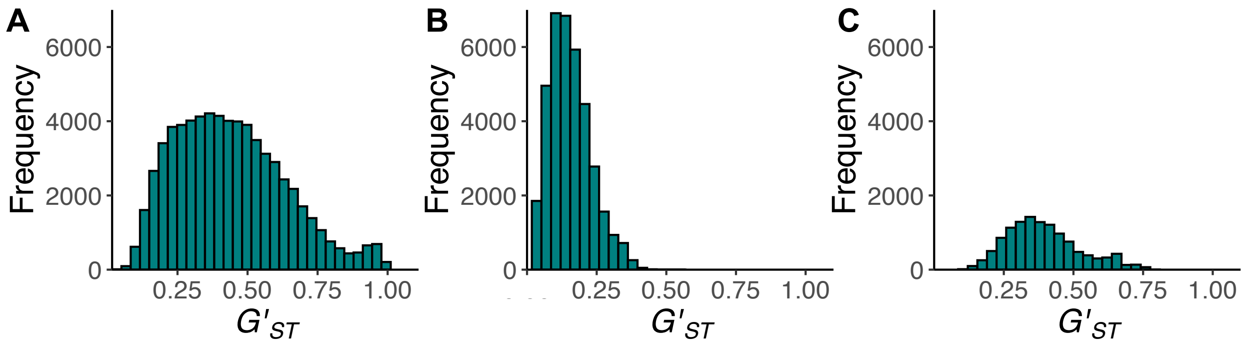


**Supplementary Fig. S6.** ***Macrophomina*** **spp. genetic variance in different subpopulations.** Fixation index distributions by **A** genotype cluster, **B** country of origin and **C** isolation source are shown. Histograms were generated with calculated $G'_{ST}$ values from the R library vcfR using Mp11-12 SNPs.

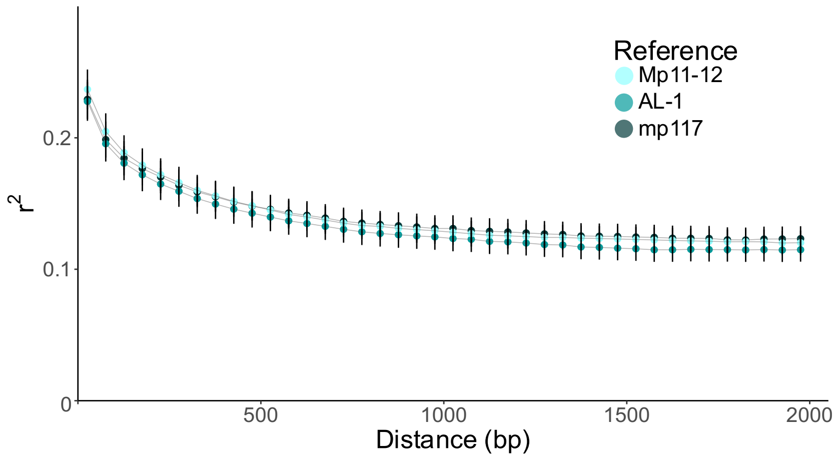


**Supplementary Fig. S7. Linkage associations of SNP loci.** Squared correlations calculated with SNPs based on *M. phaseolina* references Mp11-12, AL-1 and mp117 are shown. Error bars represent one standard error above and below the means.

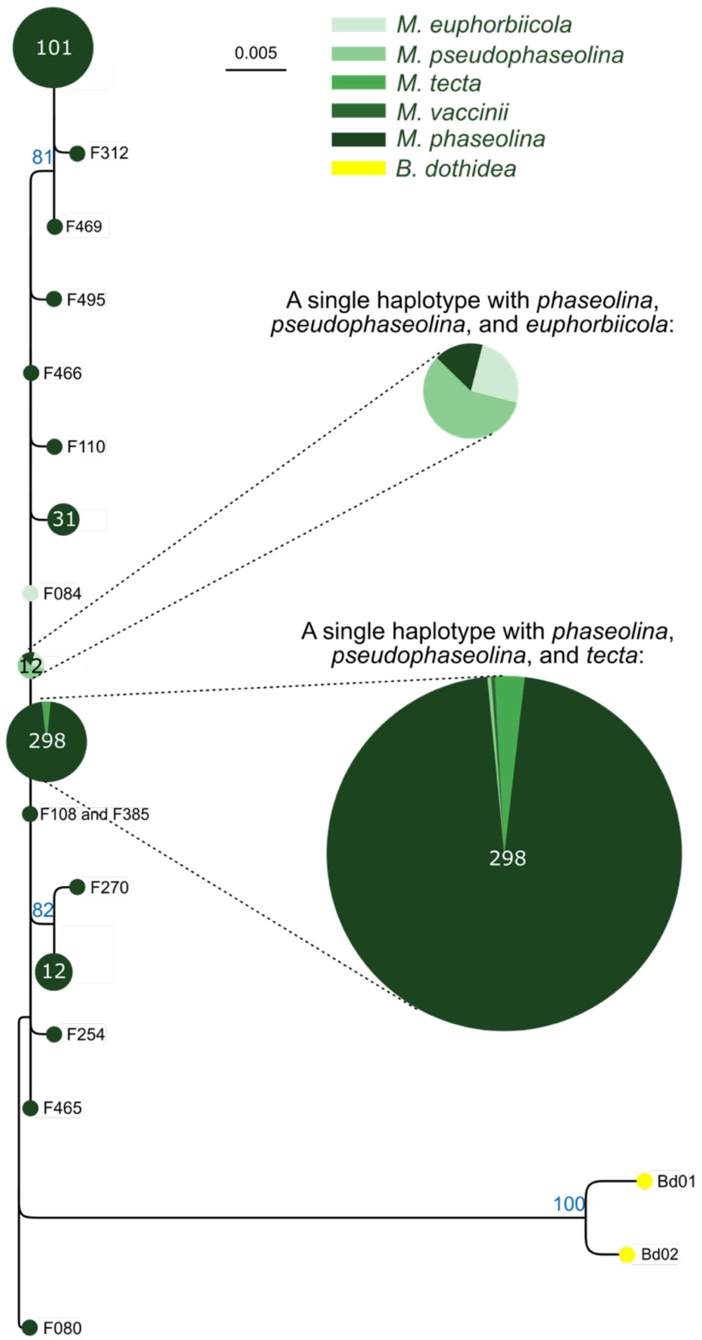


**Supplementary Fig. S8. Sequences similarities of the ITS regions of *Macrophomina* spp. isolates.** The phylogenetic tree was calculated by RAxML-NG version 1.2.1 with the GTR+G model of evolution and 1,000 bootstrap replicates. Only bootstrap values greater than 70 are shown. *Botryosphaeria dothidea* served as outgroups. Circles are approximately scaled to the number of isolates represented, which is also written in white text inside the circle. Circles representing isolates from more than one species are depicted as pie charts, with color codes shown in the upper right of the panel. Where a haplotype represents only one isolate, the isolate ID is shown to the right of the circle. The ITS region did not differentiate *Macrophomina* species, as most isolates had identical sequences. Used sequences are in Supplementary Information S1.

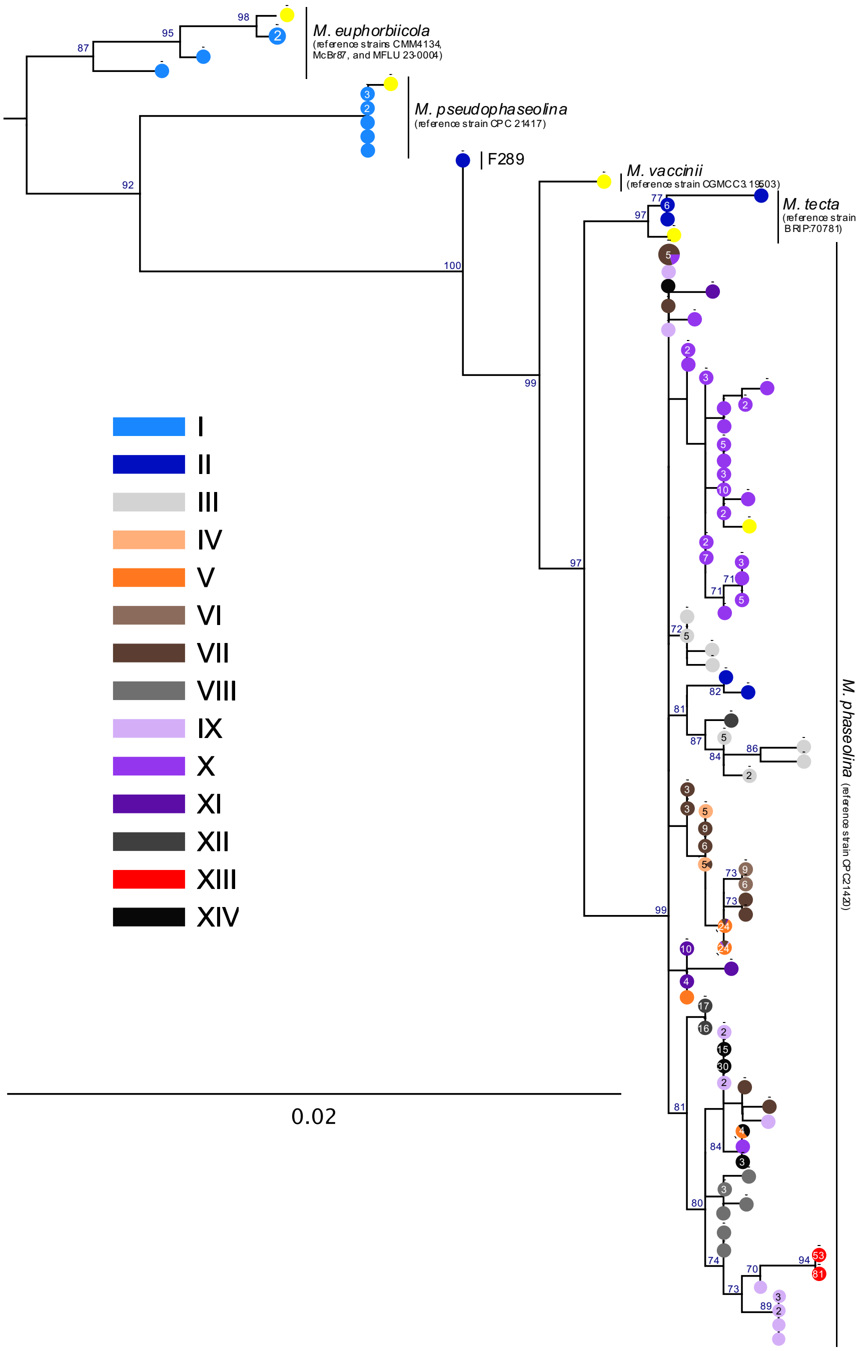


**Supplementary Fig. S9. Maximum likelihood phylogenetic tree constructed from concatenated ITS, *actin*, *beta-tubulin* and *TEF1-α* regions of *Macrophomina* spp. isolates.** The phylogenetic tree was calculated by RAxML-NG version 1.2.1 with the GTR+G model of evolution and 1,000 bootstrap replicates. Only bootstrap values greater than 70 are shown. Sequences from reference *Macrophomina* spp. genomes are denoted with yellow circles on branch tips. *Macrophomina* species could be differentiated by these combined sequences. Used sequences are in Supplementary Information S2.

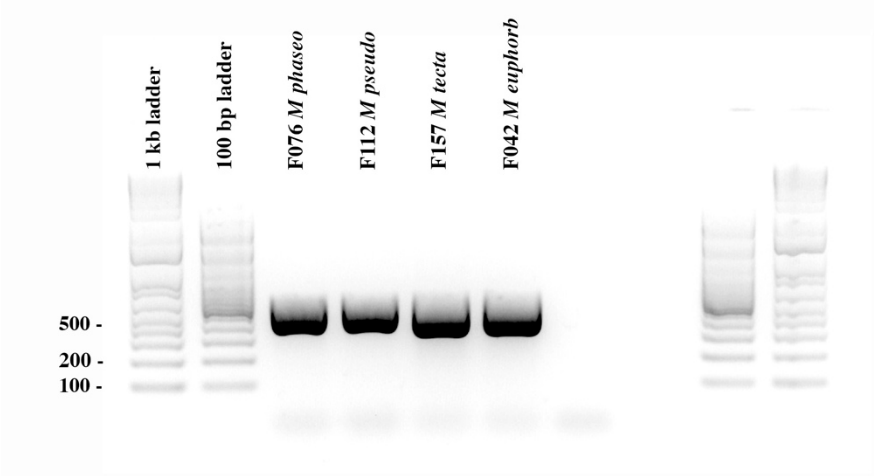


**Supplementary Fig. S10. Gel showing 'mspp' target PCR amplicons from representatives four *Macrophomina* spp.**
